# ChpC controls twitching motility-mediated expansion of *Pseudomonas aeruginosa* biofilms in response to serum albumin, mucin and oligopeptides

**DOI:** 10.1101/825596

**Authors:** Laura M. Nolan, Laura C. McCaughey, Jessica Merjane, Lynne Turnbull, Cynthia B. Whitchurch

## Abstract

Twitching motility-mediated biofilm expansion occurs via coordinated, multi-cellular collective behaviour to allow bacteria to actively expand across surfaces. Type-IV pili (T4P) are cell-associated virulence factors which mediate this expansion via rounds of extension, surface attachment and retraction. The Chp chemosensory system is thought to respond to environmental signals to regulate the biogenesis, assembly and twitching motility function of T4P. In other well characterised chemosensory systems, methyl-accepting chemotaxis proteins (MCPs) feed environmental signals through a CheW adapter protein to the histidine kinase CheA to modulate motility. The *Pseudomonas aeruginosa* Chp system has two CheW adapter proteins, PilI and ChpC, and an MCP PilJ that likely interacts via PilI with the histidine kinase ChpA. It is thought that ChpC associates with other MCPs to feed environmental signals into the system, however no such signals have been identified. In the current study we show that ChpC is involved in the response to host-derived signals serum albumin, mucin and oligopeptides. We demonstrate that these signals stimulate an increase in twitching motility, as well as in levels of 3’-5’-cyclic adenosine monophosphate (cAMP) and surface-assembled T4P. Interestingly, our data shows that changes in cAMP and surface piliation levels are independent of ChpC but that the twitching motility response to these environmental signals requires ChpC. Based upon our data we propose a model whereby ChpC associates with an MCP other than PilJ to feed these environmental signals through the Chp system to control twitching motility. The MCP PilJ and the CheW adapter PilI then modulate T4P surface levels to allow the cell to continue to undergo twitching motility. Our study is the first to link environmental signals to the Chp chemosensory system and refines our understanding of how this system controls twitching motility-mediated biofilm expansion in *P. aeruginosa*.

## Introduction

*Pseudomonas aeruginosa* is a Gram-negative WHO categorised ‘priority pathogen’ due to its high levels of antibiotic resistance and health-care associated infections. In the lungs of cystic fibrosis (CF) patients this pathogen forms biofilms resulting in chronic lung infections that are the major cause of morbidity and mortality in these individuals (1). *P. aeruginosa* is also commonly associated with chronic catheter-associated urinary tract infections (CAUTIs) (2). The success of *P. aeruginosa* as an opportunistic pathogen is largely attributed to its ability to form biofilms (3) and to produce many cell-associated and secreted virulence factors (4). Type IV pili (T4P) are a major cell-associated virulence factor which are located at the pole of the cell and are involved in biofilm formation and twitching motility-mediated active biofilm expansion via rounds of extension, surface attachment and retraction (5,6). Twitching motility is likely to facilitate active biofilm expansion by *P. aeruginosa* along the length of indwelling catheters (7–10).

The biogenesis, assembly and twitching motility function of T4P is regulated by a number of complex regulatory systems including a putative chemosensory system, the Chp system (5). This system is encoded by the *pilGHIJK*-*chpABC* gene cluster (11–14), which is homologous to the *Escherichia coli* Che chemosensory system involved in regulating flagella-mediated swimming chemotaxis in response to environmental conditions (15). The core signalling components of the Chp system in *P. aeruginosa* include a putative CheA histidine kinase homolog, ChpA (14), which appears to be coupled to the methyl-accepting chemotaxis (MCP) protein PilJ (12), via the CheW-like adapter protein, PilI (12,14). The Chp system gene cluster also encodes another CheW homolog, ChpC, which is thought to link additional MCPs to ChpA (14). *P. aeruginosa* has 26 MCPs (16,17), many of which have been linked to the sensing of environmental conditions. However, none of these MCPs, including the Chp system-associated MCP PilJ, have been shown to modulate twitching motility in response to specific external stimuli. The periplasmic domain of PilJ has however been shown to sense conformational changes in PilA, the T4P monomer unit, when the pilus is undergoing active extension, surface attachment and retraction (18). Furthermore, PilJ has been shown to interact with FimS (19), which is part of the two-component sensor FimS/AlgR that is involved in expression of *fimU-pilVWXY1Y2E* required for T4P biogenesis and assembly (5,20).

In addition to its role in controlling twitching motility, the Chp system also positively regulates intracellular levels of the second messenger, 3’-5’-cyclic adenosine monophosphate (cAMP) via the major adenylate cyclase, CyaB, which synthesises cAMP (21). The global regulator Vfr binds cAMP and activates expression of a number of genes required for T4P biogenesis (22,23). The ability of PilJ to sense retracted T4P monomer has been shown to be required for modulation of intracellular cAMP levels (18).

FimL has also been linked to the Chp system-cAMP-T4P regulatory pathway in *P. aeruginosa*. Our previous work demonstrated that this protein is necessary for twitching motility, and while *fimL* mutants have wildtype levels of intracellular T4P, surface associated T4P levels are decreased (24). In addition to the role of FimL in twitching motility, it has also been shown to involved in autolysis (24). More recent work by us and others has revealed that compensatory mutations in the cAMP phosphodiesterase *cpdA* (which breaks down cAMP) results in increased cAMP levels, restoring twitching motility in *fimL* mutants (25,26). Since this compensatory mutation in *cpdA* restores the twitching motility phenotype, FimL is likely to directly target CyaB activity, and not be involved in T4P biogenesis or assembly.

We have previously investigated the twitching motility response of *P. aeruginosa* to serum albumin (in the form of BSA), mucin and oligopeptides (in the form of tryptone) and shown that FimX is involved in the response to mucin and oligopeptides (27). *P. aeruginosa* would encounter each of these host-signals in an infection setting: mucin, which is known to be elevated in a CF lung environment (28,29), oligopeptides in urine (30–32) when *P. aeruginosa* infects a catheter-implanted urinary tract, and serum albumin in the vicinity of epithelial cells. The stimulation of twitching motility in response to these signals is likely to accelerate colonisation by *P. aeruginosa* of epithelial cells, CF lungs or implanted devices. The stimulatory response to these signals is not completely abolished in a *fimX* mutant (27) suggesting that there are additional regulators involved. In the current study we demonstrate that ChpC of the Chp chemosensory system is also involved in the twitching motility response to serum albumin, mucin and oligopeptides. Overall this study advances our understanding of how the Chp system controls T4P biogenesis and twitching motility in response to environmental signals.

## Materials and Methods

### Bacterial strains and media

The strains and plasmids used in this study and their relevant characteristics are listed in Table 1. *P. aeruginosa* was cultured on lysogeny broth (LB) solidified with agar at 1.5% (w/v) or 1% (w/v) (for interstitial biofilm expansion assays) and grown overnight at 37 °C. *P. aeruginosa* or *E. coli* cultures were grown in either cation-adjusted Mueller Hinton broth (CAMHB) or LB and incubated overnight at 37 °C, with shaking at 250 rpm. Agar plates for interstitial biofilm expansion assays consisted of base media (0.5% (w/v) yeast extract (Oxoid), 0.05% (w/v) NaCl (Sigma)) solidified with 1% (w/v) bacteriological agar (Oxoid) with or without supplements BSA (0.1% (w/v); Research Organics, USA), mucin from porcine stomach (0.05% (w/v); Sigma), tryptone (3% (w/v); Oxoid), GlcNac (50 mM) and casamino acids (3% (w/v)). Antibiotics were used at the following concentrations as required: ampicillin 50 μg/ml and tetracycline 5 μg/ml for *E. coli* and carbenicillin 250 μg/ml and tetracycline 200 μg/ml for *P. aeruginosa. P. aeruginosa* was also grown on Vogel-Bonner media (VBM) (10x solution contains MgSO4.7H2O (8  mM), citric acid (anhydrous) (9.6  mM), K2HPO4 (1.7  mM), NaNH5PO4.4H2O (22.7  mM), pH 7, and filter sterilized) with 1.5% (w/v) agar containing tetracycline (200 μg/ml) for allelic exchange mutagenesis. Counter selection and curing of pFlp2 was achieved on LB agar with 5% (w/v) sucrose. The PAK genome (NCBI number accession number LR657304; (33)) was used to search for methyl-accepting chemotaxis protein (MCP) and solute-binding protein (SBP) orthologs. The Pseudomonas.com website was also used to identify *P. aeruginosa* PA14/PAO1 orthologs (34)

**Table 1.** Strains and plasmids used in this study

| Strain or plasmid | Relevant description | Reference or source |
| --- | --- | --- |
| ***P. aeruginosa* strains** |  |  |
| PA14 | Wild type clinical *P. aeruginosa* strain | (55) |
| PAK | Wild type *P. aeruginosa* strain | D. Bradley, Memorial University of Newfoundland, St Johns, Canada |
| PA14Δ*chpC* | In frame deletion of *chpC* in wildtype strain PA14 | This study |
| PAKΔ*chpC* | In frame deletion of *chpC* in wildtype strain PAK | This study |
| PAK*pilA*:TcR | *pilA* inactivated by allelic displacement with a tetracycline resistance cassette (TetR) | (56) |
| ***E. coli* strains** |  |  |
| DH5α | *recA*, *endA1*, *gyrA96*, *hsdR17*, *thi-1*, *supE44*, *relA1*, ϕ80, *dlacZ*ΔM15 | Laboratory collection |
| S17-1 | *thi pro hsdR recA chr::RP4-2* | (57) |
| **Plasmids** |  |  |
| pUCPSK | AmpR; *E. coli*-*P. aeruginosa* shuttle vector | (56) |
| pUCP*chpC* | AmpR; A 740bp PCR amplicon containing *chpC* inserted against *Plac* into *EcoRI* site of pUCPSK | This study |
| pFlp2 | AmpR; encodes FLP recombinase and *sacB*. | (58) |
| pGEMTeasy | AmpR; *E. coli* cloning vector | Promega |
| pGemCLF | AmpR; A 1149bp PCR amplicon from the 5' end of *chpC* (CLF) and inserted into pGEMTeasy | This study |
| pGemCRF | AmpR; A 1238bp fragment PCR amplicon from the 3' end of *chpC* (CRF) and inserted into pGEMTeasy | This study |
| pCRFV | KanR; *Bam*HI-*Kpn*I fragment from pGEMCRF inserted into pOK12 | This study |
| pOKCDEL | Kan, TcR; The concatamerized fragment including the *Bgl*II-*Kpn*I fragment from pGEMCLF and the 1638bp *Kpn*I fragment from pLM1 with a *frt* flanked TcR cassette, inserted into the *Kpn*I and *Bam*HI sites of pCRFV fragment; | This study |
| pRIC380 | AmpR; *P. aeruginosa* suicide vector | (56) |
| pCDEL | AmpR, TcR; A 4008bp *Spe*I fragment from pOKCDEL containing the *chpC* deletion construct inserted into the *Spe*I site in pRIC380 against *sacB* | This study |

### Recombinant DNA Techniques

The preparation of plasmid DNA (Qiagen, Valencia, CA), restriction endonuclease digestion (New England Biolabs, Ipswich, MA), and ligation reactions (Promega, Madison, WI, and New England Biolabs) were carried out using standard protocols (35). Oligonucleotides used in the current study are listed in Table 2. The complementation plasmid pUCP*chpC* was generated by amplification of *chpC* with primers chpCF and chpcR and cloning the amplicon via pGEMT*easy* into pUCPKS. The preparation of *E. coli* competent cells and transformations were performed as previously described (35). *P. aeruginosa* competent cells were prepared by MgCl_2_ treatment and transformed as previously described (36). *P. aeruginosa* cells were prepared by sucrose treatment for electroporation and electroporated as previously described (26).

**Table 2.**
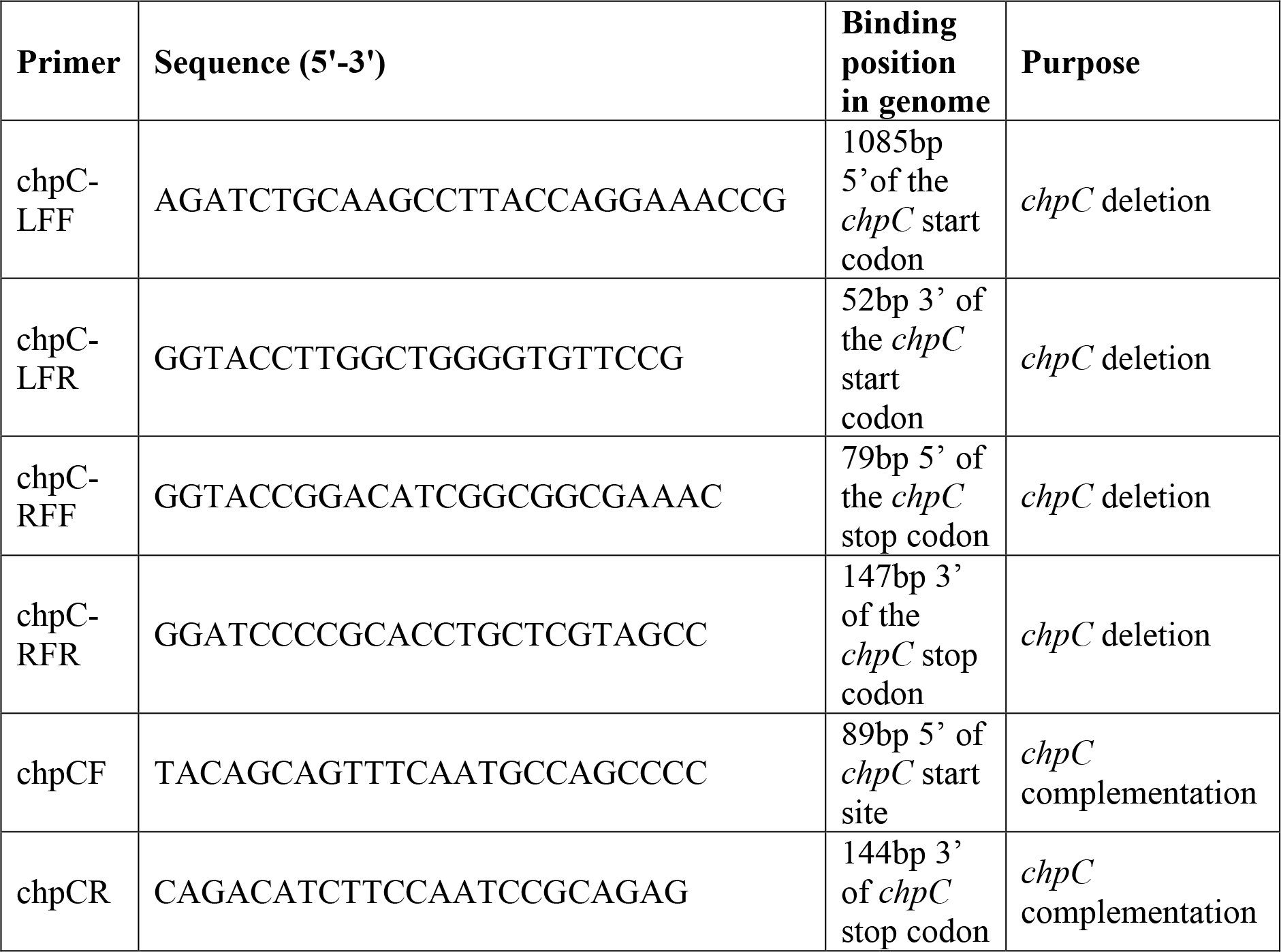
Oligonucleotides used in the current study.

### Deletion mutagenesis

To generate the in-frame *chpC* deletion construct, 1kb of sequence on both the 5’ flank of *chpC* (CLF) and the 3’ flank of *chpC* (CRF) were amplified by PCR with primer pairs chpCLFF / chpCLFR (fragment CLF) and chpCRFF/ chpCRFR (fragment CRF) from PAO1 genomic DNA. The fragment CRF was inserted into pOK12 resulting in the construct pCRFV. The fragment CLF was then inserted into pCRFV to generate pOKCDEL. The *chpC* deletion construct fragment was then sub-cloned from pOKCDEL into the suicide vector pRIC380 to generate pCDEL. *chpC* in-frame deletion mutants were made in *P. aeruginosa* strains PAK and PA14 by allelic exchange recombination followed by FRT-mediated deletion of the resistance cassette using previously published methods (37,38).

### Phenotypic assays

Interstitial biofilm expansion was assayed using a modification of the subsurface twitching motility stab assay described previously (39). Briefly, the *P. aeruginosa* strain to be tested was stab inoculated through an agar plate and cultured for 24 h at 37 °C. The longest (a) and shortest (b) diameters of each interstitial biofilm at the agar and petri dish interface were measured and the surface area calculated using the formula: area = abπ. Intracellular cAMP assays were conducted as described previously (26) using cells grown on base media supplemented with BSA (0.1% (w/v)), mucin (0.05% (w/v)) or tryptone (3% (w/v)).

### Growth experiments

Growth of *P. aeruginosa* strains was followed by recording changes in OD_595nm_ over a 20 hr period. Cells were grown in microtitre plates, and incubated at 37 °C, shaking at 250 rpm. Base media supplemented with GlcNAc (50 mM), BSA (0.1% (w/v)), mucin (0.05% (w/v)) or tryptone (3% (w/v)) was used in growth assays.

### PilA immunoblotting

Detection of cell-associated pilin was performed as described previously (14) with cells being harvested from plates grown at 37 °C for 20 hr on base media or base media supplemented with BSA (0.1% (w/v)), mucin (0.05% (w/v)) and tryptone (3% (w/v)).

### PilA ELISA

Enzyme-linked immunosorbent assays (ELISAs) were performed as described previously (14) with cells harvested from plates grown at 37 °C for 20 hr on base media or base media supplemented with BSA (0.1% (w/v)), mucin (0.05% (w/v)) or tryptone (3% (w/v)). ELISAs of cells harvested from tryptone plates were treated with 100 Kunitz units/mL DNaseI (D5025, Sigma Aldrich) for 1 hr statically at 37 °C, then washed three times with PBS, prior to use in the assay.

## Results

### ChpC is involved in the twitching motility response of *P. aeruginosa* to BSA, mucin and tryptone

BSA, mucin and tryptone have previously been shown to stimulate *P. aeruginosa* twitching motility-mediated interstitial biofilm expansion with the effect of mucin and tryptone, but not BSA, being regulated to some extent by FimX (27). However, as some stimulation of a *fimX* mutant is still observed, this suggests that other components must also be involved in mediating the twitching motility response to these host-derived signals. The *P. aeruginosa* Chp system is a putative chemosensory system that may regulate twitching motility in response to environmental signals through MCPs that intersect into the system via the second CheW homolog, ChpC (14). We were therefore interested in investigating the role of ChpC in controlling twitching motility in response to BSA, mucin and tryptone, to determine if this pathway plays a role in controlling twitching motility in response to these environmental cues.

To investigate the role of ChpC in stimulating twitching motility in response to BSA, mucin or tryptone, we first generated in-frame deletion mutants of *chpC* in *P. aeruginosa* strains PAK and PA14 and transformed the wildtype and isogenic Δ*chpC* strains with the *chpC* complementation palsmid (pUCP*chpC*) and empty vector control (pUCPSK). Interstitial biofilm expansion assays were then performed with these strains in base media agar or base media agar supplemented with BSA, mucin or tryptone. In both PAK and PA14, twitching motility-mediated interstitial biofilm expansion was significantly increased in the presence of BSA, mucin and tryptone compared to base media (Figure 1A-B). This corresponded to a relative change in the area of the interstitial biofilm for PAK of approx. 5-fold for BSA and mucin and approx. 8-fold for tryptone compared to base media (Figure 1A), whereas due to the impact of larger interstitial biofilms on base media, PA14 showed smaller relative changes in the area of interstitial biofilm of approx. 2-fold for BSA, mucin and tryptone compared to base media (Figure 1B). Interestingly, while there was still some stimulation of twitching motility for PAKΔ*chpC* on BSA, mucin and tryptone compared to base media (approx. 3-fold on BSA and mucin and approx. 4-fold on tryptone (Figure 1A)), there was no stimulation of PA14Δ*chpC* on BSA, mucin or tryptone compared to based media (Figure 1B). In both PAK and PA14 the mutation in *chpC* was fully complemented by expression of ChpC *in trans* (Figure 1A-B). To determine whether the observed stimulation of twitching motility-mediated biofilm expansion in these strains on BSA, mucin and tryptone was due to an increase in growth rate, growth assays were conducted in base media, or base media supplemented with BSA, mucin and tryptone. These growth curves revealed that BSA, mucin and tryptone did not affect the growth of the wildtype or its isogenic Δ*chpC* strain in either PAK or PA14 (Supplementary Figure 1).

**Figure 1.**
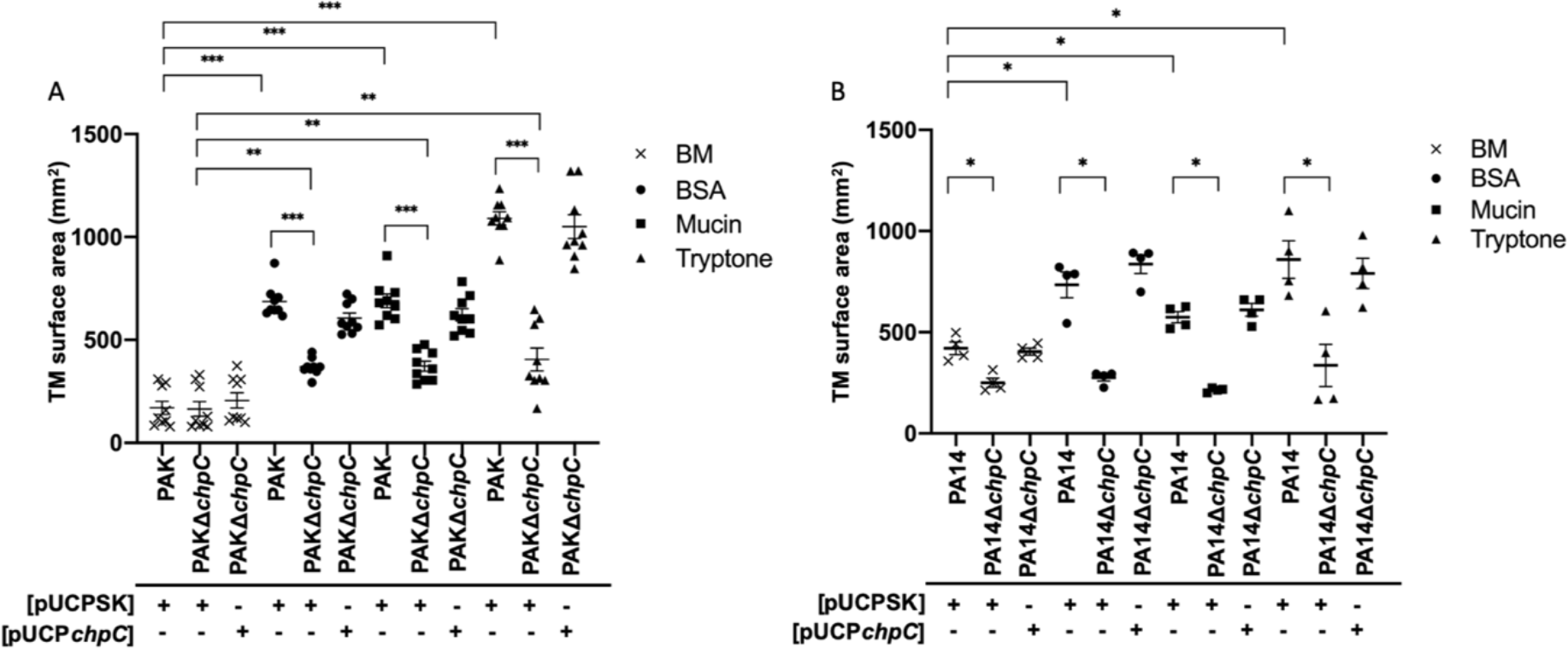
ChpC is involved in the twitching motility response of *P. aeruginosa* to BSA, mucin and tryptone. Subsurface twitching motility (TM)-mediated biofilm expansion of PAK and PAKΔ*chpC* (A) or PA14 and PA14Δ*chpC* (B) containing pUCPSK or pUCP*chpC* at the agar/plastic interstitial space after 24 hrs at 37 ̊C in base media (BM), or BM supplemented with BSA (0.1%), mucin (0.05%) or tryptone (3%). The mean of each set of technical triplicates was calculated to give an n=9 for PAK and n=4 for PA14, which is presented as mean ± SEM (* *P*<.05 *\*\*P*<.005 *\*\*\*P*<.0001; Mann-Whitney *U*-test comparing wildtype [pUCPSK] or Δ*chpC* [pUCPSK] on BM to supplemented media and wildtype [pUCPSK] to Δ*chpC* [pUCPSK] on each supplemented media. Ns for wildtype [pUCPSK] compared to Δ*chpC* [pUCP*chpC*] and ns for PA14Δ*chpC* [pUCPSK] on BM compared to supplemented media.

These results demonstrate that ChpC contributes to the twitching motility response of *P. aeruginosa* to BSA, mucin and tryptone to varying extents depending on the strain background. We went on to further characterise how *P. aeruginosa* responds to these environmental signals.

### The effect of BSA, mucin and tryptone on PilA and cAMP levels

Twitching motility levels can be affected by changes in expression of the T4P monomer subunit, PilA, and/or alterations in the rates of biogenesis/retraction of surface-assembled T4P. To determine if the mechanism of ChpC-mediated stimulation of twitching motility in response to BSA, mucin and tryptone involved changes in expression and/or assembly of the T4P, we performed ELISAs to measure levels of surface-assembled T4P and immunoblots to measure levels of the T4P subunit PilA in whole cells harvested from confluent lawns of PAK and PAKΔ*chpC* grown on base media, or base media supplemented with BSA, mucin or tryptone. The ELISAs demonstrated that both PAK and PAKΔ*chpC* grown on BSA and mucin had marginally increased levels of surface-assembled T4P compared to when grown on base media (Figure 2A, Supplementary Figure 2). We observed a significant amount of autolysis in the confluent lawns of cells cultured on tryptone as has been previously reported (24) and ELISAs of cells harvested directly from the tryptone plates were unable to detect any surface PilA in either strain, which suggested that there may be excessive amounts of extracellular DNA (eDNA) present. To remove this eDNA, cells were harvested from both base media and tryptone plates, treated with DNaseI and then washed in PBS, prior to use in the ELISA. This allowed detection of surface-assembled T4P in these cells and revealed that both PAK and PAKΔ*chpC* cells grown in the presence of tryptone had 2 to 3-fold higher levels of surface-assembled T4P than cells cultured on base media (Figure 2A, Supplementary Figure 2). Whole cell immunoblots demonstrated that there were no changes in PilA expression levels for wildtype or PAKΔ*chpC* when grown on base media, or base media supplemented with BSA, mucin or tryptone (Figure 2B).

**Figure 2.**
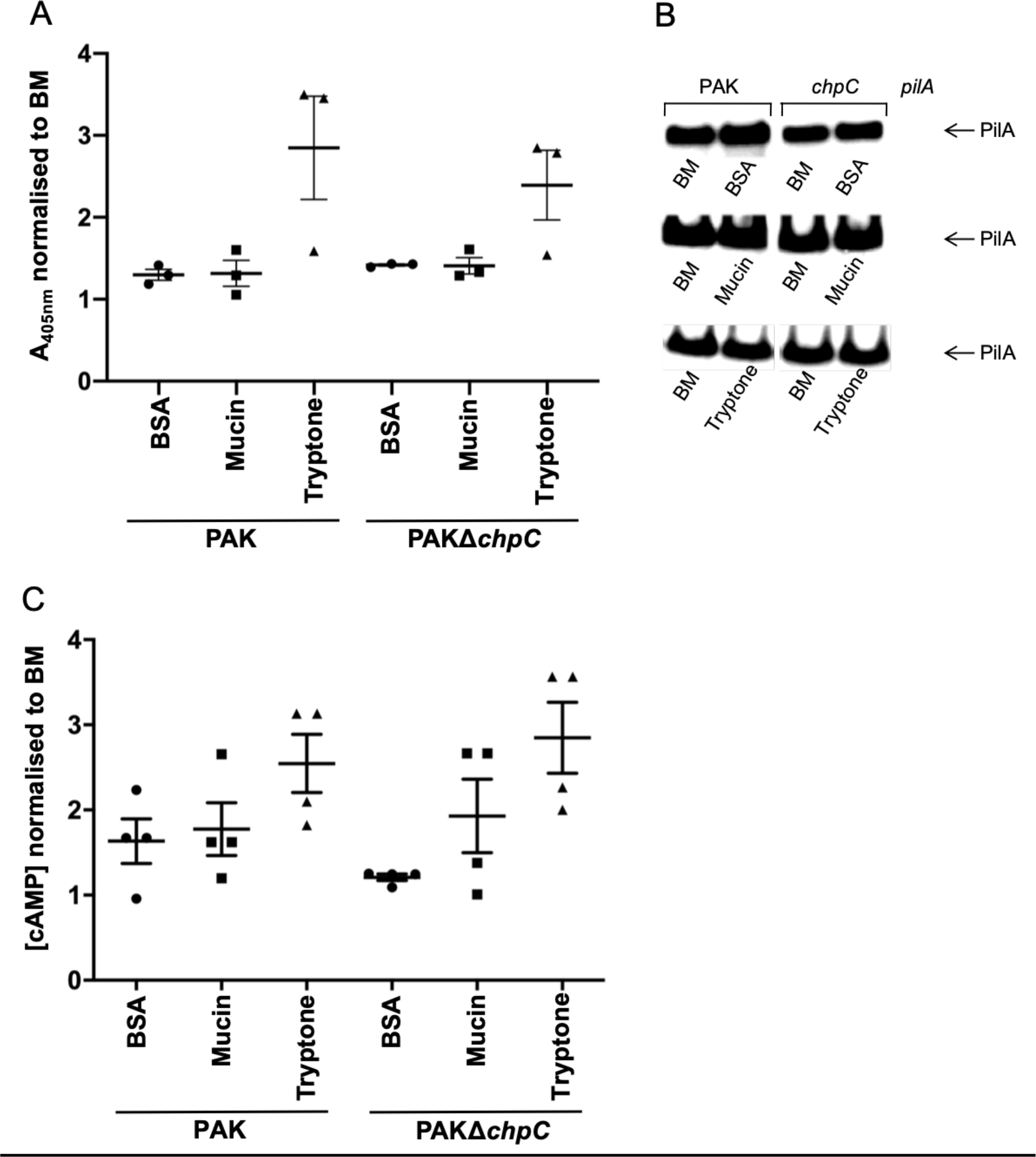
The effect of BSA, mucin and tryptone on PilA and cAMP levels. (A) PilA ELISAs of wildtype PAK and PAKΔ*chpC* grown on base media (BM) or BM supplemented with BSA (0.1%), mucin or tryptone (3%). This data is based upon the raw data shown in Supplementary Figure 2. Here the mean of technical triplicates for the most concentrated sample for PAK or PAKΔ*chpC* on BM for each biological replicate (n=3) was used to normalise the mean of technical triplicates of PAK and PAKΔ*chpC* on each supplement for the respective biological replicate. The data are presented as mean ± SEM. Equal cell numbers were loaded. (B) PilA Western immunoblot of whole cell lysates from wildtype PAK and PAKΔ*chpC* cells grown on BM or BM supplemented with BSA (0.1%), mucin (0.05%), tryptone (3%), and PAK*pilA* cells grown on BM. Equal cell numbers were loaded. (C) Intracellular cAMP concentrations (pmol/mL) of wildtype PAK and PAKΔ*chpC* cells grown in base media (BM) or BM supplemented with BSA (0.1%), mucin (0.05%) or tryptone (3%). The mean of technical triplicates for PAK or PAKΔ*chpC* on BM for each biological replicate (n=4) was used to normalise the mean of technical triplicates of PAK and PAKΔ*chpC* on each supplement for the respective biological replicate. The data are presented as mean ± SEM. Equal cell numbers were loaded.

The *P. aeruginosa* Chp system has been shown to positively regulate cAMP levels via CyaB to increase levels of surface-assembled T4P (21). To investigate the involvement of cAMP in the twitching motility response of *P. aeruginosa* to BSA, mucin and tryptone, intracellular cAMP levels were measured in cells harvested from confluent lawns of PAK and PAKΔ*chpC* grown on base media, or base media supplemented with BSA, mucin or tryptone. For both PAK and PAKΔ*chpC* cAMP levels were marginally increased on BSA and mucin and 2 to 3-fold on tryptone (Figure 2C), which reflects the changes in surface-assembled pili with these supplements (Figure 2A and Supplementary Figure 2).

These observations suggest that stimulating twitching motility results in higher levels of cAMP and surface-assembled T4P in both PAK and PAKΔ*chpC* although there does not appear to be a direct correlation between these levels and rates of interstitial biofilm expansion. This is particularly evident in PAKΔ*chpC* which showed equivalent increases in interstitial biofilm sizes in BSA, mucin and tryptone (Figure 1A) but much greater increases in the levels of cAMP and surface-assembled T4P in tryptone than in BSA or mucin (Figure 2 and Supplementary Figure 2). Thus, it appears that changes in cAMP and surface piliation levels are largely independent of ChpC but that stimulation of biofilm expansion in response to these environmental cues is modulated via ChpC.

### Examination of potential chemical ligands that stimulate twitching motility

Chemosensory systems typically sense, via their MCP components, small chemical ligands such as amino acids, inorganic phosphate and chlorinated and non-chlorinated hydrocarbons (17). Given this, we wanted to determine if the twitching motility responses to BSA, mucin and tryptone were mediated by recognition of a smaller component common to each of these host-derived signals. BSA consists of a single polypeptide chain consisting of 583 amino acid residues, the mucin used in this study is a mixture of highly glycosylated glycoproteins obtained by pepsin digestion of hog stomach, while tryptone is the assortment of oligopeptides and a few free amino acids, formed by the digestion of casein by the protease trypsin. To determine if amino acids are the common smaller signal sensed to modulate twitching motility, we performed subsurface stab assays with wildtype PAK and PA14 strains on casamino acids, which are produced by acid hydrolysis of casein, resulting in mainly free amino acids, and a few oligopeptides. Casamino acids did not stimulate twitching motility in either strain and repressed rather than stimulated twitching motility in PAK (Figure 3A), which suggests that amino acids are not the smaller component of BSA, mucin and tryptone that is sensed by *P. aeruginosa* to stimulate twitching motility.

**Figure 3.**
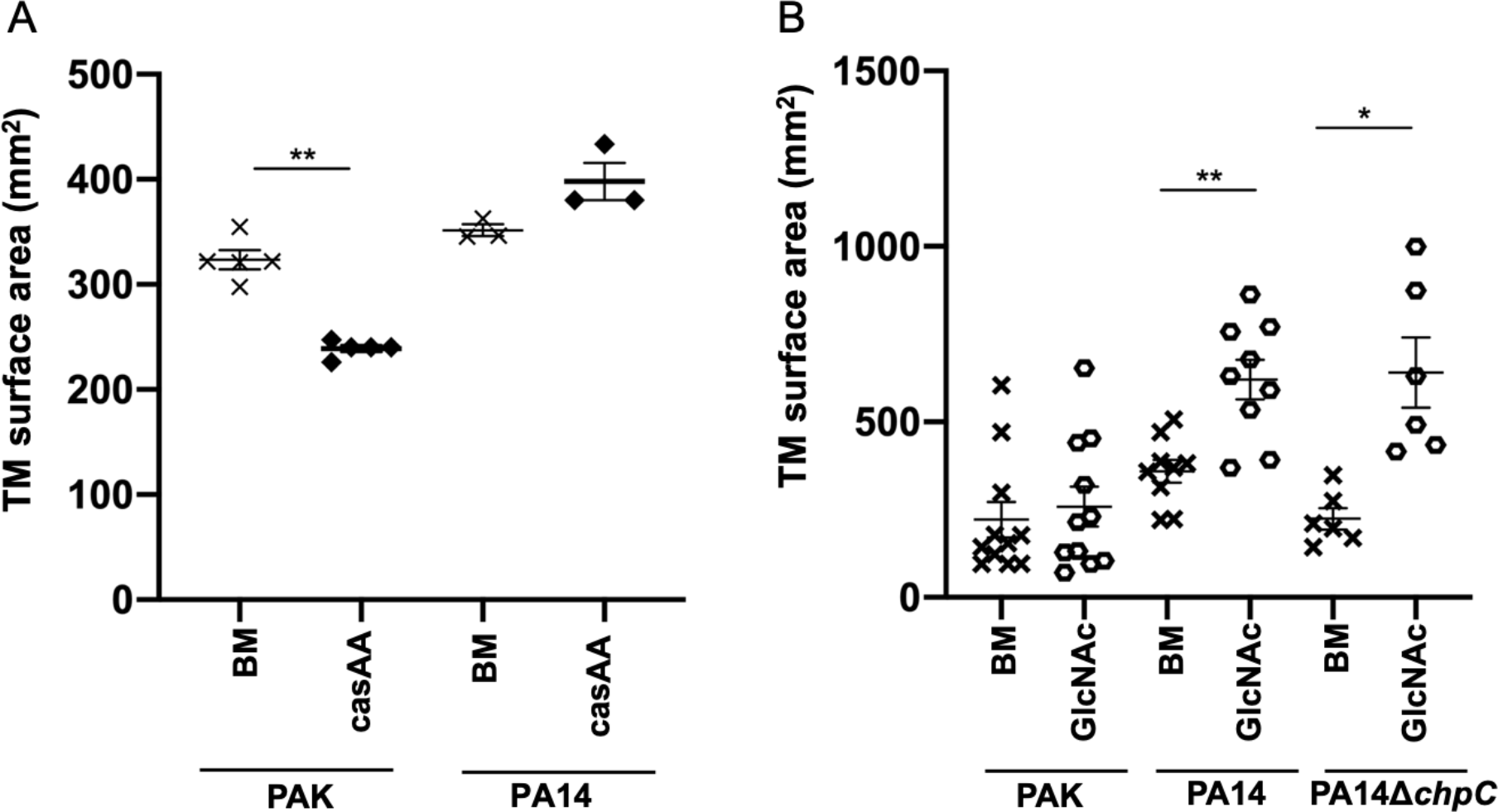
The twitching motility response of *P. aeruginosa* to casamino acids and GlcNac. Subsurface twitching motility (TM)-mediated biofilm expansion at the agar/plastic interstitial space after 24 hrs at 37 ̊C of (A) PAK or PA14 on base media (BM), or BM supplemented with casamino acids (casAA) (3%): the mean of each set of technical triplicates was calculated to give an n≥3 which is presented as mean ± SEM. (*\*\*P*<0.005; Mann-Whitney *U*-test compared to BM; PA14 BM compared to casamino acids ns *(P* = 0.100)). (B) PAK, PA14 and PA14Δ*chpC* on BM or BM supplemented with GlcNAc (50mM): the mean of each set of technical triplicates was calculated to give an n≥6 which is presented as mean ± SEM (\*\**P*<.005, *\*P*<0.05; Mann-Whitney *U*-test compared to BM, PAK BM compared to GlcNAc ns (*P* = 0.6518)).

Mucin is a highly glycosylated glycoprotein with up to 90% of its molecular weight due to O- and N-linked oligosaccharides. As N-acetylglucosamine (GlcNAc) is a common sidechain of mucin we were interested in determining if GlcNAc might stimulate *P. aeruginosa* twitching motility. As GlcNAc is present in CF sputum at mM concentrations (29) we therefore tested the twitching motility response of PAK and PA14 at a GlcNAc concentration within this range (50 mM). Whilst there was no significant increase in PAK twitching motility in GlcNAc compared to base media (Figure 3B), PA14 was significantly stimulated by GlcNAc resulting in an approx. 2-fold increase in interstitial biofilm area relative to base media (Figure 3B). Growth assays with strain PA14 in media supplemented with GlcNAc confirmed that the observed stimulation of twitching motility was not simply due to an increased growth rate, as cells grew at the same rate in base media as in base media supplemented with GlcNAc (Supplementary Figure 1E). As ChpC appears to be necessary for the twitching motility response to mucin in strain PA14 (Figure 1), we examined the twitching motility response of PA14Δ*chpC* to GlcNAc and found that PA14Δ*chpC* was stimulated by GlcNAc to the same extent as wildtype PA14 (Figure 3B). This indicates that while twitching motility of *P. aeruginosa* is stimulated by GlcNAc in strain PA14, this is not the component of mucin that is being sensed to modulate twitching motility via ChpC.

## Discussion

In the study we have investigated the twitching motility response of *P. aeruginosa* to the host-derived signals serum albumin (in the form of BSA), mucin, and oligopeptides (in the form of tryptone). We found that each of these stimulate twitching motility-mediated biofilm expansion and increase the levels of surface-assembled T4P and intracellular cAMP to varying extents. Interestingly, our observations indicate that there is not a direct relationship between the levels of surface assembled T4P, cAMP and rates of interstitial biofilm expansion. We also determined that the Chp chemosensory system appears to be involved in regulating twitching motility-mediated interstitial biofilm expansion in response to these signals and that this occurs, at least in part, via ChpC. However, stimulation of T4P and cAMP levels in response to mucin, BSA and tryptone is independent of ChpC.

In the current study we show that both *P. aeruginosa* strains PAK and PA14 are stimulated by BSA, mucin and tryptone but that the degree of this stimulation and the involvement of ChpC in this response varies (Figure 1). This is unsurprising considering the large amount of anecdotal evidence for phenotypic variation between *P. aeruginosa* laboratory strains. For PA14 it appears that the response to these signals is fed solely through the Chp system via ChpC, since there were no differences in the amount of twitching motility of PA14Δ*chpC* on base media compared to any of the supplements (Figure 1B). In contrast, we found that twitching motility in PAKΔ*chpC* is still stimulated to some extent by these signals (Figure 1A), which suggests the response is fed through the Chp system via ChpC and another component(s) in PAK. This is possibly through the other Chp system-associated CheW-adapter, PilI. However, given that mixing of MCP and CheW proteins has been shown to occur in chemosensory systems in other bacteria (40), it is also possible that another CheW homolog from one of the other four *P. aeruginosa* chemosensory systems feeds the environmental signal from the MCP to ChpA. It is also possible that in PAK some residual stimulation by mucin and tryptone occurs through FimX in the absence of ChpC.

*P. aeruginosa* possesses 26 MCPs and of these only PilJ has so far been shown to be associated with the Chp system (12). Recent studies have demonstrated that *P. aeruginosa* surface association results in an increase in cAMP levels via PilJ (19) and that this is due to PilJ sensing conformational changes in retracted PilA in the periplasm, which promotes CyaB-dependent cAMP generation, to increase expression of components required for T4P assembly (18). Our data show that BSA, mucin and tryptone cause an increase in cAMP levels, to different extents, but that in all cases this is not reliant upon the presence of ChpC (Figure 2C). This suggests that sensing of these environmental signals requires ChpC to be linked to one of the other 25 *P. aeruginosa* MCPs (16,17), and not to the cAMP modulating MCP PilJ. Furthermore, our data shows that simply increasing T4P levels is not sufficient to increase the rate of twitching motility expansion, but that ChpC is also required to direct this expansion. This is demonstrated by PAKΔ*chpC* having the same levels of twitching motility on BSA, mucin and tryptone (Figure 1A), despite having markedly higher levels of surface T4P on tryptone (Figure 2A).

We propose a model (Figure 4) in which BSA, mucin and tryptone are sensed by one or more MCPs other than PilJ, which are linked to the Chp system via ChpC to stimulate twitching motility. The degree to which twitching motility is stimulated would then reflect the amount of retracted T4P in the periplasm, *i.e* cells on tryptone undergo more twitching motility, which would result in an increase in levels of retracted T4P in the periplasm. This is then sensed by the MCP PilJ, which modulates cAMP levels to control expression of T4P components accordingly, in order to supply the required amount of pilin subunit for the biogenesis of surface T4P so the cell can continue to undergo twitching motility.

**Figure 4.**
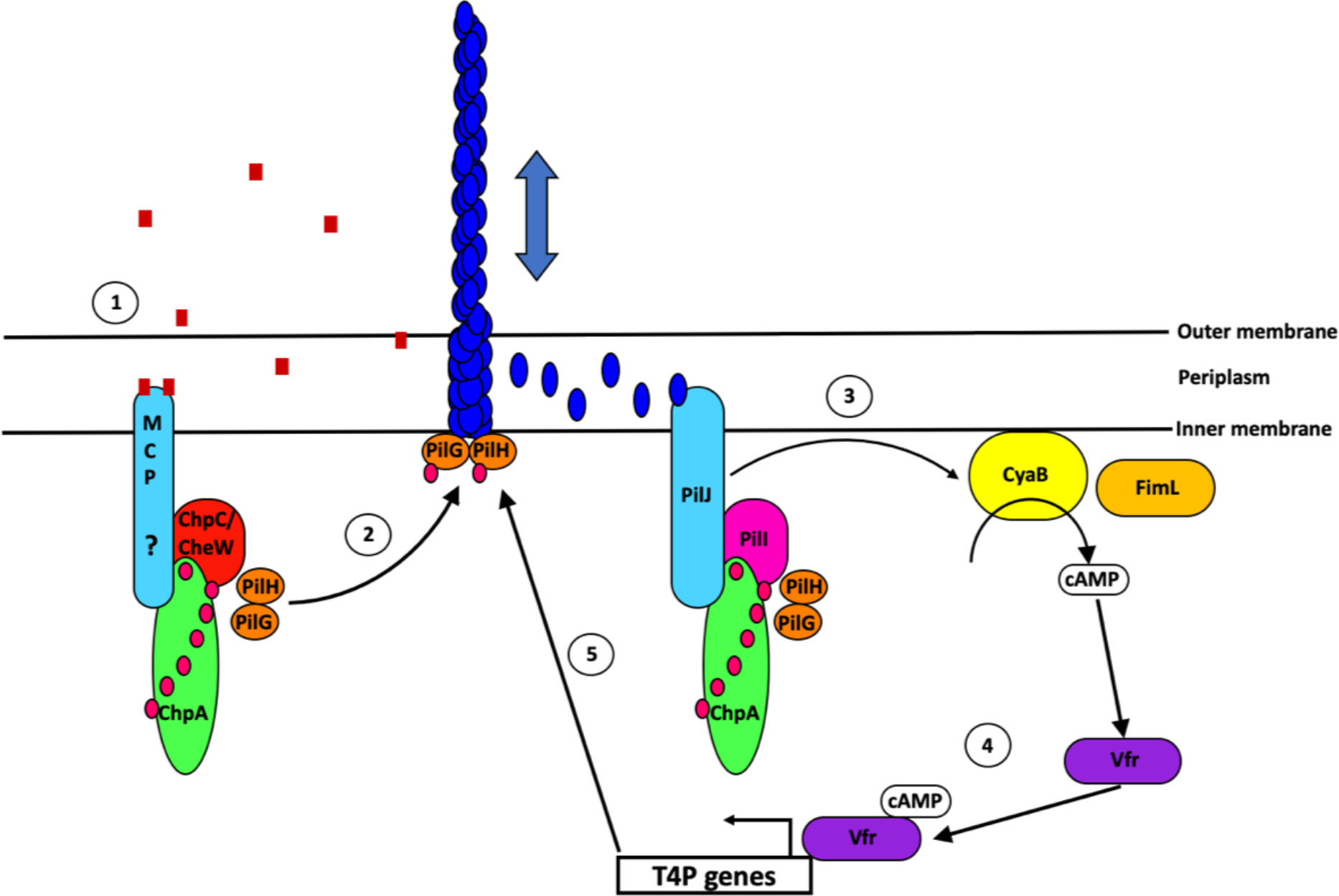
Chp system sensing of environmental signals and modulation of twitching motility and T4P levels. 1) Environmental signal (red squares) is sensed by an MCP which transfers the signal via ChpC/other CheW homolog to ChpA. 2) A twitching motility response is mediated by extension and retraction of the T4P. 3) PilJ senses the levels of retracted T4P in the periplasm and stimulates the generation of cAMP via CyaB. FimL affects CyaB activity. 4) cAMP-Vfr activates expression of T4P components. 5) These increase levels of assembled T4P to allow the cell to continue to undergo twitching motility in response to the environmental signal.

Since chemosensory systems will typically sense small chemical ligands via the MCP component, we also investigated whether any potential degradation products of BSA, mucin and tryptone stimulate twitching motility. We found that twitching motility is not stimulated by casamino acids (Figure 3A). Since both tryptone and casamino acids are derived from casein - the former a mixture of oligopetides and the latter a mixture of amino acids - our observations suggest that the signal sensed via ChpC is likely to be oligopeptides. For BSA and mucin it is possible that one or more of the large number of secreted proteases (41) is required for degradation of these proteins to generate oligopeptides. Indeed, previous reports have demonstrated that the twitching motility response of *P. aeruginosa* to phosphatidylcholine (PC) requires degradation of PC by the phospholipase C PlcB to release diacylglycerol (DAG). The long-chain fatty acid (LCFA) moiety of DAG is then involved in directing twitching motility of *P. aeruginosa* up PC gradients (42).

In this study we observed a considerable amount of autolysis on the confluent lawn plates of *P. aeruginosa* grown on tryptone (as we have previously reported (24)). We have previously shown that eDNA facilitates the efficient expansion of *P. aeruginosa* biofilms (6). Thus, it is possible that the additional eDNA available to tryptone-grown cells may further stimulate twitching motility and explain the marked increase in overall twitching motility of *P. aeruginosa* on tryptone (Figure 1).

Since GlcNAc is the major sidechain of mucin we also investigated if this was able to stimulate twitching motility. Interestingly, we observed that twitching motility of PA14, but not PAK was stimulated by GlcNAc. This response was not mediated by ChpC (Figure 3B). Therefore, the ChpC-mediated response of PA14 and PAK to mucin must be due to sensing of a different smaller component of mucin, such as oligopeptides or another of the monosaccharides commonly found in mucins such as galactose, fucose, N-acetylgalactosamine (GalNAc) or sialic acid.

We hypothesise that the strain differences we observed between PAK and PA14 may be due to variations in the complement of MCPs in these strains. However, this does not appear to be the case as both strains have orthologues of all 26 *P. aeruginosa* MCPs. Chemotaxis in *E. coli* can also involve a solute binding protein (SBP) that interacts with an MCP to mediate chemotaxis. A recent study has identified 98 putative SBPs in *P. aeruginosa* PAO1 (43). We compared the complement of these SBPs in the annotated genomes of PAO1, PAK and PA14 and found that one putative PAO1 SBP, PA0203, is absent in both PA14 and PAK. Another point of difference between PAO1, PA14 and PAK putative SBPs is with PA0288 (LapA)/PA0289 (LapB). PAK has an essentially identical (99% amino acid identity) orthologue of the putative PAO1 LapA (PA0688), and an orthologue with reasonable identity (68% amino acid identity) to LapB (PA0689), both of which are absent in PA14. However, as has been previously identified, PA14 does not possess LapA or LapB but has another orthologue, LapC, which has 45% amino acid identity to LapA and 40% amino acid identity to LapC. These proteins are thought to be part of a phosphate scavenging or sensing system required in low phosphate environments (44). So, whilst some differences exist in the repertoire of the LapA/B/C putative SBPs in PA14 and PAK these are unlikely to explain the observed differences in GlcNAc sensing. This suggests that there may an additional SBP or MCP in PA14, or that variations may exist in the ligand binding domains of MCPs and/or SBPs between PA14 and PAK, or that under the conditions of our assay the expression of the components responsible for sensing vary in PAK and PA14.

*P. aeruginosa* is commonly associated with a range of infections including those in the lungs of CF individuals (45), ulcerative keratitis (46), otitis externa (47), infections of the skin and soft-tissue (48), as well as in hospital-acquired infections, such as pneumonia (49) and CAUTIs (50). *P. aeruginosa* would encounter serum albumin in all of these environments, due to the proximity to different epithelial cells. Within the lungs of CF patients, *P. aeruginosa* would also encounter relatively high levels of mucin and GlcNac which are known to be elevated in the CF lung (28,29) and in CAUTIs *P. aeruginosa* would encounter oligopeptides present in urine (32). The stimulation of twitching motility in response to these signals is likely to accelerate surface colonisation by *P. aeruginosa* of epithelial cells or implanted devices within an infected host.

GlcNAc-induced stimulation of *P. aeruginosa* twitching motility is also likely to be relevant within polymicrobial environments. Gram-positive bacteria are known to shed peptidoglycan, which contains GlcNAc in a polymeric form, in large quantities (51,52). The ability of *P. aeruginosa* to uptake and sense GlcNAc has respective roles in mediating lysis of *Staphylococcus aureus* and *Bacillus licheniformis*, and in stimulating production of the virulence factors elastase and pyocyanin (29,53). Stimulation of twitching motility by GlcNAc may provide a competitive advantage within a polymicrobial setting by allowing *P. aeruginosa* to navigate closer to the source of GlcNAc, for instance a Gram-positive bacterial cell, providing more effective delivery of virulence factors.

Overall, the observations presented here add to our understanding of how *P. aeruginosa* responds to a number of relevant host-derived signals to modulate twitching motility. As *P. aeruginosa* possesses a large number of regulatory systems that intersect to control the biogenesis and function of T4P and twitching motility (54), it is clear that the regulation of twitching motility is indeed complex. However, in most cases, it has not yet been determined which signals modulate the various regulatory pathways to co-ordinate twitching motility. While this adds an additional level of complexity, this will allow us to develop a more complete understanding of how twitching motility is regulated in infection and environmental settings.

## Authors and contributions

L.M.N., L.C.M., J.M. and C.B.W. conducted experiments. L.M.N and C.B.W. analysed results. C.B.W. and L.T. provided project supervision. C.B.W. provided project administration and funding. L.M.N. and C.B.W. wrote the manuscript.

## Acknowledgements

The authors would like to acknowledge Dervilla McGowan who performed some experimental work leading up to the work presented in this manuscript.

---

**Supplementary Figure 1.**
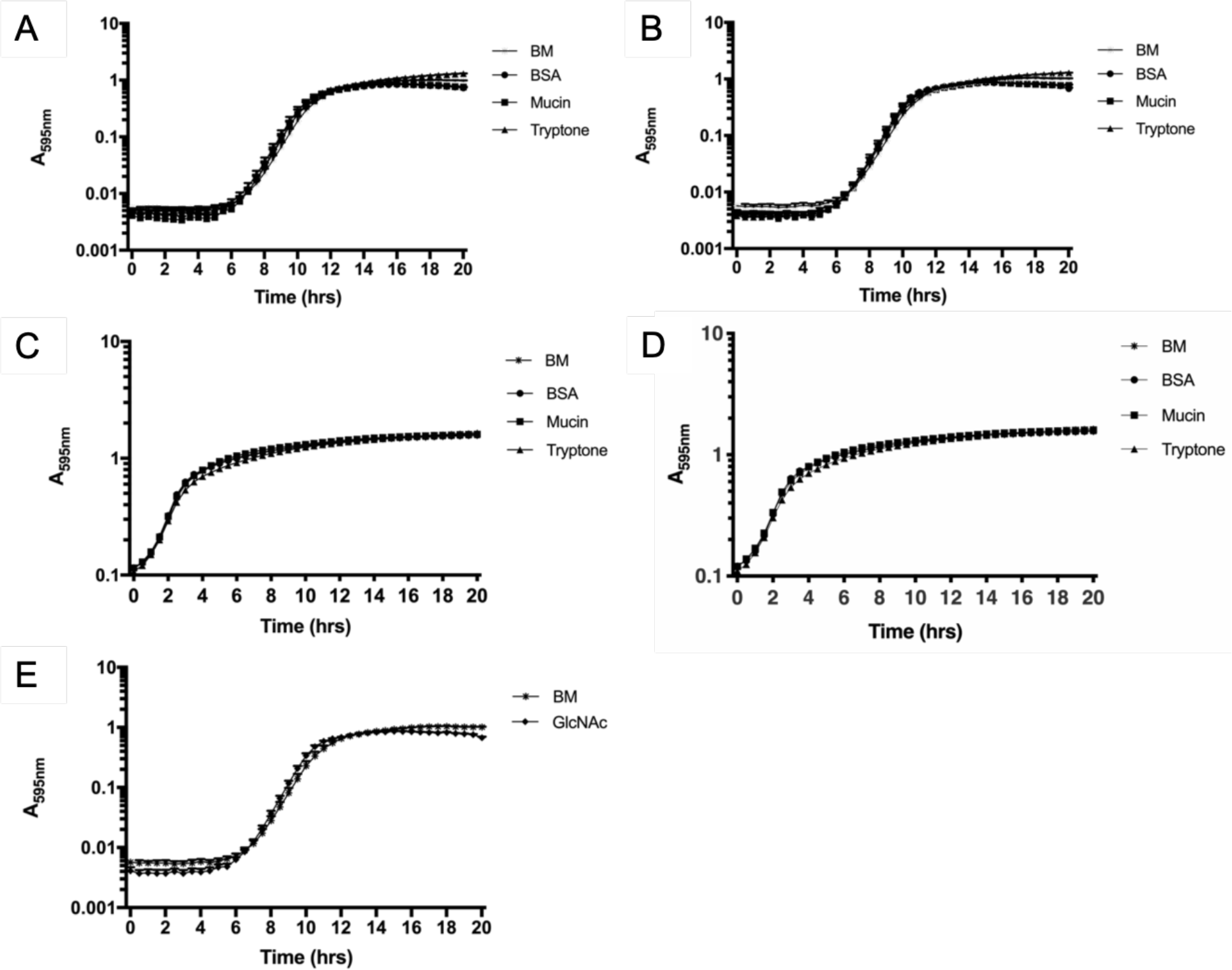
Growth rates of PAK and PA14 do not account for increased twitching motility on supplements. Growth assays over 20 hrs at 37 in base media (BM), or BM supplemented with BSA (0.1%), mucin (0.05%), tryptone (3%) or GlcNAc (50 mM) for PAK (A), PAKΔ*chpC* (B), PA14 (C, E) and PA14Δ*chpC* (D). The mean ± SEM for 3 independent experiments is presented.

**Supplementary Figure 2.**
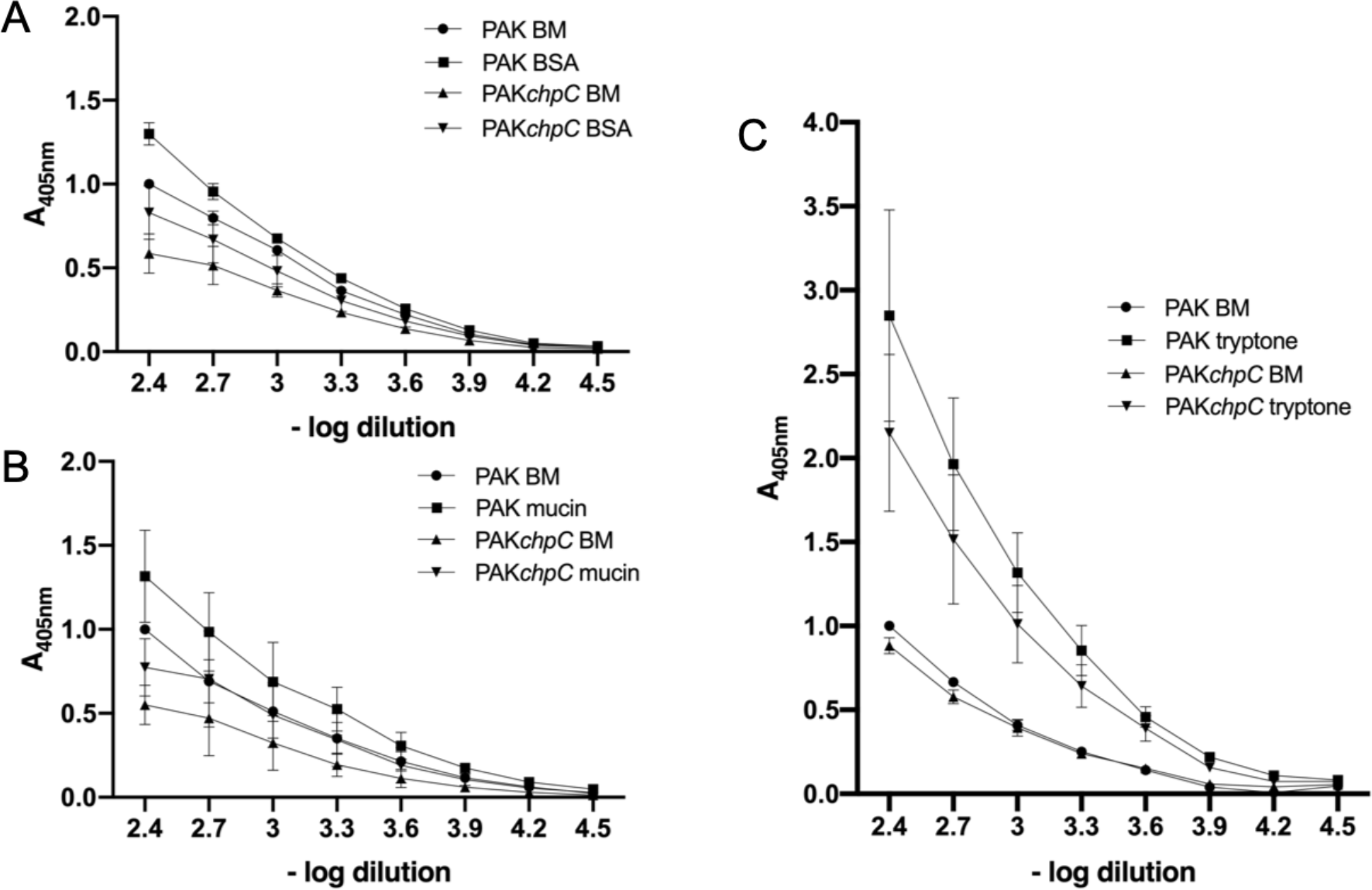
The effect of BSA, mucin and tryptone on PilA surface assembly. PilA ELISAs of wildtype PAK, PAKΔ*chpC* cells grown on base media (BM) or BM supplemented with (A) BSA (0.1%), (B) mucin (0.05%) or (C) tryptone (3%). The mean of each set of technical triplicates was calculated for PAK on BM at the lowest dilution (−log 2.4). This was then used to normalise the mean of each set of technical triplicates for PAK on supplemented BM or PAKΔ*chpC* on BM or supplemented BM for a total of 3 biological replicates. The data are presented as mean ± SEM. Equal cell numbers were loaded.

## References

1. Høiby N, Ciofu O, Bjarnsholt T. Pseudomonas aeruginosa biofilms in cystic fibrosis. Future Microbiol. 2010 Nov;5(11):1663–74.

2. Mittal R, Aggarwal S, Sharma S, Chhibber S, Harjai K. Urinary tract infections caused by *Pseudomonas aeruginosa*: a minireview. J Infect Public Health. 2009 Sep 19;2(3):101–11.

3. Donlan RM, Costerton JW. Biofilms: survival mechanisms of clinically relevant microorganisms. Clin Microbiol Rev. 2002 Apr;15(2):167–93.

4. Van Delden C, Iglewski BH. Cell-to-cell signaling and *Pseudomonas aeruginosa* infections. Emerging Infect Dis. 1998 Dec;4(4):551–60.

5. Whitchurch CB. Biogenesis and function of type IV pili in pseudomonas species. In: Ramos J-L, Levesque RC, editors. Pseudomonas. Springer US; 2006. p. 139–88.

6. Gloag ES, Turnbull L, Huang A, Vallotton P, Wang H, Nolan LM, et al. Self-organization of bacterial biofilms is facilitated by extracellular DNA. Proc Natl Acad Sci USA. 2013 Jul 9;110(28):11541–6.

7. Donlan RM. Biofilms and device-associated infections. Emerging Infect Dis. 2001 Apr;7(2):277–81.

8. Sabbuba N, Hughes G, Stickler DJ. The migration of *Proteus mirabilis* and other urinary tract pathogens over Foley catheters. BJU Int. 2002 Jan;89(1):55–60.

9. Stickler DJ. Bacterial biofilms in patients with indwelling urinary catheters. Nat Clin Pract Urol. 2008 Nov;5(11):598–608.

10. Nickel JC, Downey J, Costerton JW. Movement of pseudomonas aeruginosa along catheter surfaces. A mechanism in pathogenesis of catheter-associated infection. Urology. 1992 Jan;39(1):93–8.

11. Darzins A. The pilG gene product, required for *Pseudomonas aeruginosa* pilus production and twitching motility, is homologous to the enteric, single-domain response regulator CheY. J Bacteriol. 1993;175(18):5934–44.

12. Darzins A. Characterization of a *Pseudomonas aeruginosa* gene cluster involved in pilus biosynthesis and twitching motility: sequence similarity to the chemotaxis proteins of enterics and the gliding bacterium *Myxococcus xanthus*. Mol Microbiol. 1994;11(1):137–53.

13. Darzins A. The pilG gene product, required for *Pseudomonas aeruginosa* pilus production and twitching motility, is homologous to the enteric, single-domain response regulator CheY. Mol Microbiol. 1995;15(4):703–17.

14. Whitchurch CB, Leech AJ, Young MD, Kennedy D, Sargent JL, Bertrand JJ, et al. Characterization of a complex chemosensory signal transduction system which controls twitching motility in *Pseudomonas aeruginosa*. Mol Microbiol. 2004 May;52(3):873–93.

15. Baker MD, Wolanin PM, Stock JB. Signal transduction in bacterial chemotaxis. Bioessays. 2006 Jan;28(1):9–22.

16. Croft L, Beatson SA, Whitchurch CB, Huang B, Blakeley RL, Mattick JS. An interactive web-based Pseudomonas aeruginosa genome database: discovery of new genes, pathways and structures. Microbiology (Reading, Engl). 2000 Oct;146 (Pt 10):2351–64.

17. Kato J, Kim HE, Takiguchi N, Kuroda A, Ohtake H. *Pseudomonas aeruginosa* as a model microorganism for investigation of chemotactic behaviors in ecosystem. J Biosci Bioeng. 2008 Jul;106(1):1–7.

18. Persat A, Inclan YF, Engel JN, Stone HA, Gitai Z. Type IV pili mechanochemically regulate virulence factors in *Pseudomonas aeruginosa*. Proc Natl Acad Sci USA. 2015 Jun 16;112(24):7563–8.

19. Luo Y, Zhao K, Baker AE, Kuchma SL, Coggan KA, Wolfgang MC, et al. A hierarchical cascade of second messengers regulates *Pseudomonas aeruginosa* surface behaviors. MBio. 2015 Jan 27;6(1).

20. Belete B, Lu H, Wozniak DJ. *Pseudomonas aeruginosa* AlgR regulates type IV pilus biosynthesis by activating transcription of the fimU-pilVWXY1Y2E operon. J Bacteriol. 2008 Mar;190(6):2023–30.

21. Fulcher NB, Holliday PM, Klem E, Cann MJ, Wolfgang MC. The *Pseudomonas aeruginosa* Chp chemosensory system regulates intracellular cAMP levels by modulating adenylate cyclase activity. Mol Microbiol. 2010 May;76(4):889–904.

22. Wolfgang MC, Lee VT, Gilmore ME, Lory S. Coordinate regulation of bacterial virulence genes by a novel adenylate cyclase-dependent signaling pathway. Dev Cell. 2003 Feb;4(2):253–63.

23. Beatson SA, Whitchurch CB, Sargent JL, Levesque RC, Mattick JS. Differential regulation of twitching motility and elastase production by Vfr in Pseudomonas aeruginosa. J Bacteriol. 2002 Jul;184(13):3605–13.

24. Whitchurch CB, Beatson SA, Comolli JC, Jakobsen T, Sargent JL, Bertrand JJ, et al. Pseudomonas aeruginosa fimL regulates multiple virulence functions by intersecting with Vfr-modulated pathways. Mol Microbiol. 2005 Mar;55(5):1357–78.

25. Inclan YF, Huseby MJ, Engel JN. FimL regulates cAMP synthesis in Pseudomonas aeruginosa. PLoS One. 2011 Jan 11;6(1):e15867.

26. Nolan LM, Beatson SA, Croft L, Jones PM, George AM, Mattick JS, et al. Extragenic suppressor mutations that restore twitching motility to *fimL* mutants of *Pseudomonas aeruginosa* are associated with elevated intracellular cyclic AMP levels. Microbiologyopen. 2012 Dec;1(4):490–501.

27. Huang B, Whitchurch CB, Mattick JS. FimX, a multidomain protein connecting environmental signals to twitching motility in Pseudomonas aeruginosa. J Bacteriol. 2003 Dec;185(24):7068–76.

28. Li JD, Dohrman AF, Gallup M, Miyata S, Gum JR, Kim YS, et al. Transcriptional activation of mucin by *Pseudomonas aeruginosa* lipopolysaccharide in the pathogenesis of cystic fibrosis lung disease. Proc Natl Acad Sci U S A. 1997;94(3):967–72.

29. Korgaonkar AK, Whiteley M. Pseudomonas aeruginosa enhances production of an antimicrobial in response to N-acetylglucosamine and peptidoglycan. J Bacteriol. 2011 Feb;193(4):909–17.

30. Putnam DF. Composition and concentrative properties of human urine. NASA contract report. 1971;July.

31. Lutz W, Markiewicz K, Klyszejko-Stefanowicz L. Oligopeptides excreted in the urine of healthy humans and of patients with nephrotic syndrome. Clin Chim Acta. 1972 Jul;39(2):425–31.

32. Kentsis A, Monigatti F, Dorff K, Campagne F, Bachur R, Steen H. Urine proteomics for profiling of human disease using high accuracy mass spectrometry. Proteomics Clin Appl. 2009 Sep 1;3(9):1052–61.

33. Cain AK, Nolan LM, Sullivan GJ, Whitchurch CB, Filloux A, Parkhill J. Complete Genome Sequence of Pseudomonas aeruginosa Reference Strain PAK. Microbiol Resour Announc. 2019 Oct 10;8(41).

34. Winsor GL, Griffiths EJ, Lo R, Dhillon BK, Shay JA, Brinkman FSL. Enhanced annotations and features for comparing thousands of Pseudomonas genomes in the Pseudomonas genome database. Nucleic Acids Res. 2016 Jan 4;44(D1):D646–53.

35. Sambrook J RD. Molecular Cloning: A Laboratory Manual. Cold Spring Harbor, NY: Cold Spring Harbor Laboratory Press; 2001.

36. Mattick JS, Bills MM, Anderson BJ, Dalrymple B, Mott MR, Egerton JR. Morphogenetic expression of. J Bacteriol. 1987;169(1):33–41.

37. Hoang TT, Kutchma AJ, Becher A, Schweizer HP. Integration-proficient plasmids for *Pseudomonas aeruginosa*: site-specific integration and use for engineering of reporter and expression strains. Plasmid. 2000 Jan;43(1):59–72.

38. Choi K-H, Gaynor JB, White KG, Lopez C, Bosio CM, Karkhoff-Schweizer RR, et al. A Tn7-based broad-range bacterial cloning and expression system. Nat Methods. 2005 Jun;2(6):443–8.

39. Semmler AB, Whitchurch CB, Mattick JS. A re-examination of twitching motility in *Pseudomonas aeruginosa*. Microbiology (Reading, Engl). 1999;145:2863–73.

40. O’Neal L, Gullett JM, Aksenova A, Hubler A, Briegel A, Ortega D, et al. Distinct Chemotaxis Protein Paralogs Assemble into Chemoreceptor Signaling Arrays To Coordinate Signaling Output. MBio. 2019 Sep 24;10(5).

41. Hoge R, Pelzer A, Rosenau F, Wilhelm S. Weapons of a pathogen: proteases and their role in virulence of *Pseudomonas aeruginosa*. Current Research, Technology and Education Topics in Applied Microbiology and Microbial Biotechnology. 2010. p. 383–95.

42. Barker AP, Vasil AI, Filloux A, Ball G, Wilderman PJ, Vasil ML. A novel extracellular phospholipase C of *Pseudomonas aeruginosa* is required for phospholipid chemotaxis. Mol Microbiol. 2004 Aug;53(4):1089–98.

43. Fernández, Rico-Jiménez, Ortega, Daddaoua, García García, Martín-Mora, et al. Determination of Ligand Profiles for Pseudomonas aeruginosa Solute Binding Proteins. ijms. 2019 Oct 17;20(20):5156.

44. Ball G, Viarre V, Garvis S, Voulhoux R, Filloux A. Type II-dependent secretion of a Pseudomonas aeruginosa DING protein. Res Microbiol. 2012 Jul 23;163(6-7):457–69.

45. Sousa AM, Pereira MO. *Pseudomonas aeruginosa* Diversification during Infection Development in Cystic Fibrosis Lungs-A Review. Pathogens. 2014 Aug 18;3(3):680–703.

46. Bourcier T, Thomas F, Borderie V, Chaumeil C, Laroche L. Bacterial keratitis: predisposing factors, clinical and microbiological review of 300 cases. Br J Ophthalmol. 2003 Jul;87(7):834–8.

47. Roland PS, Stroman DW. Microbiology of acute otitis externa. Laryngoscope. 2002 Jul;112(7 Pt 1):1166–77.

48. Lipsky BA, Tabak YP, Johannes RS, Vo L, Hyde L, Weigelt JA. Skin and soft tissue infections in hospitalised patients with diabetes: culture isolates and risk factors associated with mortality, length of stay and cost. Diabetologia. 2010 May;53(5):914–23.

49. Kollef MH, Shorr A, Tabak YP, Gupta V, Liu LZ, Johannes RS. Epidemiology and outcomes of health-care-associated pneumonia: results from a large US database of culture-positive pneumonia. Chest. 2005 Dec;128(6):3854–62.

50. Aguilar-Duran S, Horcajada JP, Sorlí L, Montero M, Salvadó M, Grau S, et al. Community-onset healthcare-related urinary tract infections: comparison with community and hospital-acquired urinary tract infections. J Infect. 2012 May;64(5):478–83.

51. Doyle RJ, Chaloupka J, Vinter V. Turnover of cell walls in microorganisms. Microbiol Rev. 1988 Dec;52(4):554–67.

52. Mauck J, Chan L, Glaser L. Turnover of the cell wall of Gram-positive bacteria. J Biol Chem. 1971 Mar 25;246(6):1820–7.

53. Korgaonkar A, Trivedi U, Rumbaugh KP, Whiteley M. Community surveillance enhances *Pseudomonas aeruginosa* virulence during polymicrobial infection. Proc Natl Acad Sci USA. 2013 Jan 15;110(3):1059–64.

54. Burrows LL. Pseudomonas aeruginosa twitching motility: type IV pili in action. Annu Rev Microbiol. 2012 Jul 2;66:493–520.

55. Rahme LG, Stevens EJ, Wolfort SF, Shao J, Tompkins RG, Ausubel FM. Common virulence factors for bacterial pathogenicity in plants and animals. Science. 1995 Jun 30;268(5219):1899–902.

56. Watson AA, Alm RA, Mattick JS. Construction of improved vectors for protein production in *Pseudomonas aeruginosa*. Gene. 1996 Jan;172(1):163–4.

57. Simon R, O’Connell M, Labes M, Pühler A. Plasmid vectors for the genetic analysis and manipulation of rhizobia and other gram-negative bacteria. Plant Molecular Biology. Elsevier; 1986. p. 640–59.

58. Hoang TT, Karkhoff-Schweizer RR, Kutchma AJ, Schweizer HP. A broad-host-range Flp-FRT recombination system for site-specific excision of chromosomally-located DNA sequences: application for isolation of unmarked *Pseudomonas aeruginosa* mutants. Gene. 1998 May 28;212(1):77–86.

